# Polyunsaturated fatty acids promote appetite via the microbiome-gut-brain axis

**DOI:** 10.1101/2025.07.29.667447

**Authors:** Yi Jia Liow, Shusei Eshima, Mustafa Talay, Vladimir Yeliseyev, Lynn Bry, Rachel N. Carmody

## Abstract

Appetite is regulated by nutrient-sensing systems that integrate long-term signals from energy stores and short-term cues from dietary intake, yet this regulation is increasingly disrupted by industrialized diets. Although the physiological effects of industrialized diets are well documented, the continued rise in metabolic and eating disorders underscores a critical gap in our understanding of how these diets shape neural regulation of eating behavior. Here, we tested how distinct properties of industrialized diets alter brain neurochemistry and change appetite. We probed the properties of an industrialized diet through contrasts targeting the overall diet pattern (Western vs. control), enriched macronutrients (fat vs. sugar), and isocaloric trade-offs of macronutrient variants (saturated fatty acids vs. polyunsaturated fatty acids [PUFA]). The most salient effects emerged from the finest-grained contrast: PUFA conditioning increased appetite through a mechanism involving elevated brain 5-hydroxyindoleacetic acid (5-HIAA), a primary serotonin catabolite associated with the gut microbiome. Fecal microbiota transplants into germ-free mice confirmed that the PUFA-conditioned gut microbiota carries an appetite-enhancing signature. Together, our findings delineate a diet-microbiome-gut-brain axis through which dietary components of industrialized diets can modulate appetite and contribute to altered eating behavior.

**Graphical abstract:** 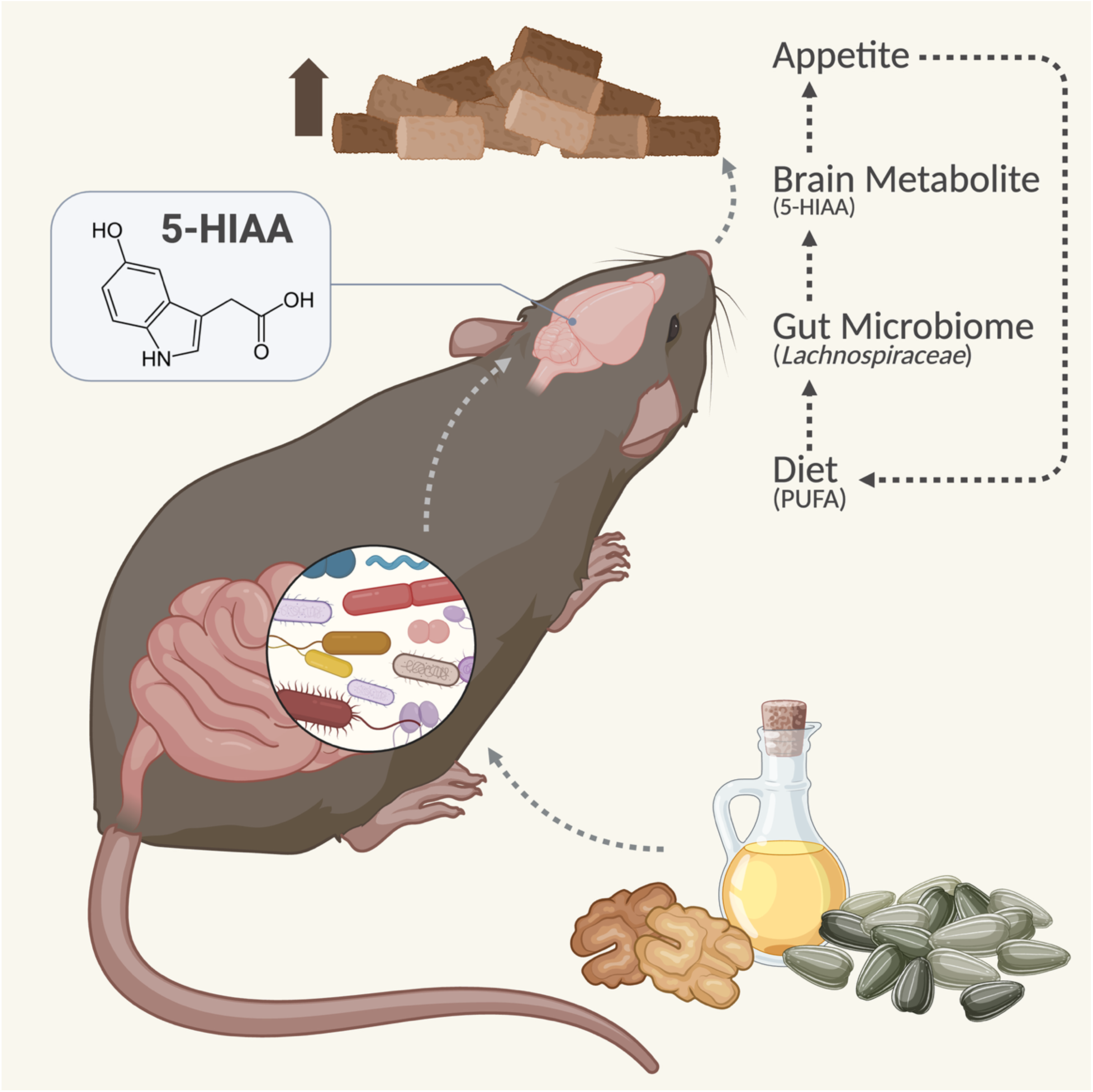

**One Sentence Summary:** Dietary polyunsaturated fatty acids enhance appetite via a gut microbiome–serotonergic pathway.

## INTRODUCTION

Appetite regulation is a tightly coordinated physiological process in which nutrient-derived signals, shaped by dietary composition, act on central circuits to regulate energy intake (*1,2*). Industrialized dietary patterns can disrupt this system by altering nutrient-derived signals in ways that overwhelm systems of metabolic control shaped by evolutionary pressures (*3,4*). The high prevalence and persistence of obesity and metabolic disorders remain difficult to treat (*5*), highlighting a gap in our understanding of how modern diets impair appetite-regulating signals.

Industrialized diets rich in refined carbohydrates and saturated fatty acids but low in fiber impair metabolic homeostasis and alter feeding behavior by remodeling satiety signaling and reward valuation (*6–8*). These nutrient profiles bias consumption toward energy-dense foods while disrupting internal signals that regulate intake (*9*). While broad contrasts between industrialized and non-industrialized diets have revealed global effects on metabolism (*10*) and eating behavior (*11*), they often obscure the specific dietary components that can distort how the brain registers the nutritional quality of food (*12*). Among other possible pathways, industrialized diets are known to alter the gut microbiome (*13*) and microbiome-derived metabolites modulate satiety via the gut-brain axis (*14–16*), suggesting a role for the microbiome-gut-brain axis in the impairment of appetite regulation (*8, 17*). However, which specific components of an industrialized diet contribute to shaping host appetite via the microbiome-gut-brain axis remains unexplored.

We hypothesized that industrialized diets perturb appetite regulation through microbe-derived metabolites that engage host metabolic and neural pathways. We thus sought to identify the pathways through which specific dietary components enriched in industrialized diets influence appetite. We narrowed the dietary components of interest through feeding trials in mice targeting the overall diet pattern (Western vs. control), enriched macronutrients (fat vs. sugar), and isocaloric trade-offs of macronutrient variants (saturated fatty acids [SFA] vs. polyunsaturated fatty acids [PUFA]). Across each dietary contrast, we probed the relationships between appetite, dietary composition, microbiome profile, and metabolites known to interact with host metabolism (*18*), dietary intake (*19–22*), and the gut microbiome (*15,18,19*). Many of these metabolites – including neurotransmitters such as gamma-aminobutyric acid (GABA) and serotonin, bile acids, short-chain fatty acids, and endocannabinoids – are bioactive, capable of signaling nutritional status to the brain and influencing feeding behavior through gut-brain communication pathways (*8, 25*). Using brain-targeted metabolomics, we aimed to identify metabolite signatures involved in appetite regulation.

The finest-grained contrast, SFA vs. PUFA, identified 5-hydroxyindoleacetic acid (5-HIAA) as a metabolite linking PUFA dietary conditioning, gut microbiome profile, and appetite. Follow-up gnotobiotic experiments confirmed that the PUFA-conditioned gut microbiome carries an appetite-promoting signature that does not depend on direct dietary exposure. Jointly, this work characterizes a diet-microbiome-gut-brain axis through which fatty acid subclasses shape brain metabolite profiles associated with appetite, providing mechanistic insight into how industrialized diets could perpetuate unhealthy eating behavior.

## RESULTS

### Dietary conditioning alters host appetite

We subjected C57BL/6J mice (n=10) to a multi-stage dietary trial involving a baseline 24-hour intake assay, followed by two weeks of exposure to a conditioning diet, followed by a post-intervention 24-hour intake assay, and finally six days of re-exposure to the conditioning diet prior to sacrifice and tissue harvest (Fig. 1A). Our conditioning diets were designed to model the effects of industrialized diets at three levels of specificity: overall dietary pattern, comparing Western versus control diet-mimic chows; enriched macronutrients, comparing high-fat versus high-sugar chows; and trade-offs of macronutrient variants, comparing isocaloric SFA-versus PUFA-rich chows (Fig. 1B, Tables S1-S4). In each 24-hour intake assay, the two contrasted conditioning diets were presented alongside each other, with total food intake and diet-specific intake measured via reduced hopper weights and validated based on continuous video recordings analyzed with DeepLabCut, a machine-learning-based software for pose estimation (*26*).

**Fig. 1:**
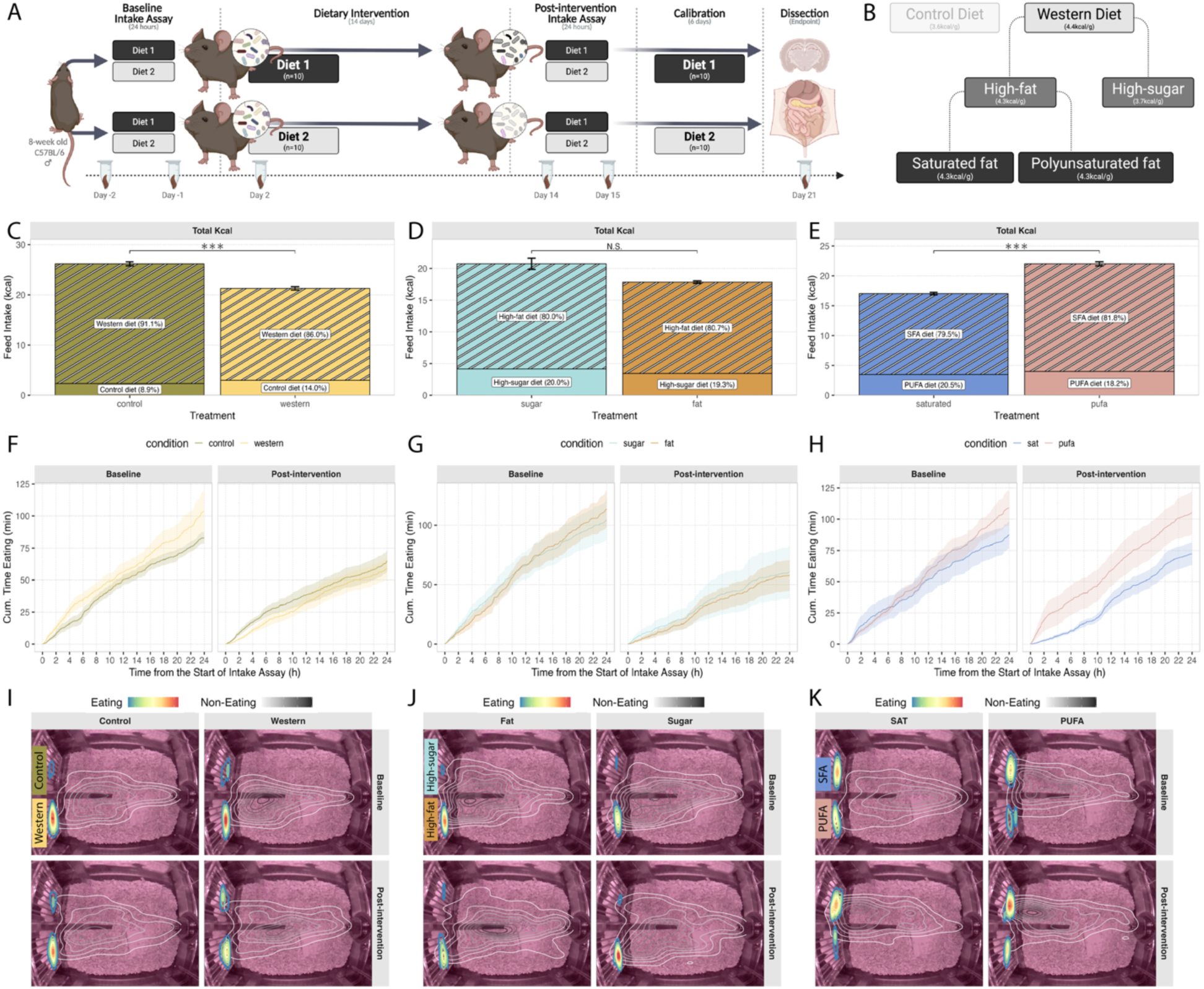
Dietary conditioning modulates appetite. **A-B** Schematics of the conventional mouse experiment (**A**) and dietary contrasts tested as nested properties of the Western diet (**B**). Images created in BioRender. **C-E** Total caloric intake during the post-intervention 24-hour intake assay for the Western vs. control (**C**), fat vs. sugar (**D**), and SFA vs. PUFA (**E**) dietary contrasts. Western, high-fat, and SFA diet conditioning suppressed total intake relative to controls (n = 10/group). Shading denotes dietary source of calories during the assay. Wilcoxon rank sum test, *\*\*\* adj-p <* 0.05, N.S. = not significant. **F-H** DeepLabCut-derived cumulative eating time across 24-hour intake assays at baseline and post-intervention; solid lines indicate group means and shaded bands denote mean ± SEM. **I-K** DeepLabCut-generated spatial heatmaps depicting average positional density of mice during the 24-hour intake assays, with color-scaled contours marking time spent eating and greyscale contours marking non-eating bouts at baseline (top row) and post-intervention (bottom row) for each dietary contrast.

We first examined how dietary conditioning influenced appetite by quantifying and comparing total caloric intake across groups in the 24-hour intake assays. Western-conditioned mice consumed fewer kilocalories than control-conditioned mice post-intervention (Fig. 1C). No difference was detected between fat- and sugar-conditioned groups, although intake trended lower in the fat-conditioned group (Fig. 1D). PUFA-conditioned mice, by contrast, consumed more kilocalories than SFA-conditioned mice (Fig. 1E).

To determine whether appetite shifts targeted specific nutrient classes, we further analyzed intake at the level of macronutrients (carbohydrate, fat, protein), carbohydrate subclasses (simple, complex, indigestible), and fatty acid subclasses (SFA, monounsaturated fatty acids [MUFA], or PUFA) (Fig S1). The Western-conditioned group exhibited lower intake than their control-conditioned counterparts, with the strongest effects observed for macronutrient fat (Fig. S1A) and SFA consumption (Fig. S1I). Unlike the Western vs. control contrast, sugar- vs. fat-conditioning did not elicit a robust shift in nutrient-specific intake (Fig. S1B, S1F, S1J). While most nutrient consumption scaled with total caloric intake (Fig. S1C, S1G), PUFA-conditioned mice consumed more SFA and MUFA than their SFA-conditioned counterparts (Fig. S1K), indicating that PUFA conditioning enhanced appetite specifically for fatty acid subclasses absent from the conditioning diet. Together, these results show that exposure to Western, high-fat, and SFA-rich diets consistently reduced total caloric intake, whereas PUFA conditioning enhanced appetite in part through elevated SFA consumption.

While endpoint intake measurements reveal how much was consumed, they offer limited insight into how consumption patterns unfold over time. To address this, we applied DeepLabCut to extract temporal features of feeding behavior from video recordings during the intake assays. We defined eating as instances when the mouse nose point remained in contact with the feeder for more than two consecutive seconds and quantified the total duration of these events over the 24-hour period as a proxy for consumption. This complementary analysis uncovered diet-specific changes in temporal patterns of feeding behavior. In the Western vs. control contrast, the control-conditioned group exhibited consumption patterns that were similar at baseline and post-intervention, whereas the Western-conditioned group showed a 43.6% decrease in cumulative eating time post-intervention (Fig. 1F). In the fat vs. sugar contrast, cumulative eating time decreased in both groups, by 49.2% and 41.7%, respectively (Fig. 1G). In the SFA vs. PUFA contrast, cumulative eating time in the PUFA-conditioned group changed minimally (−3.59%) despite an increase in caloric intake – indicating a faster eating rate – while the SFA-conditioned group exhibited a 17.5% decrease in cumulative eating time (Fig. 1H).

Heatmaps of consummatory activity further illustrate preferences between the two co-presented diets. In the Western vs. control and fat vs. sugar contrasts, all mice predominantly consumed from the Western and high-fat feeders, respectively, regardless of timepoint (Fig. 1I, 1J). However, post-intervention activity density around the feeders was attenuated in Western- and fat-conditioned mice, suggesting reduced post-intervention engagement with their assigned conditioning diets. In the SFA vs. PUFA contrast, both groups developed a clear preference for the SFA diet, as evidenced by more defined activity density around the SFA feeder post-intervention (Fig. 1K).

To provide physiological context, we assessed cumulative feed intake, percent body weight change, and feed efficiency during the two-week dietary intervention, as well as fat mass (via EchoMRI) at the start and end of this period (Fig. S2). Western-conditioned mice consumed more of their assigned diet than did their control counterparts through day 12 (Fig. S2A) and exhibited greater fat mass (Fig. S2D) despite no differences in body weight gain (Fig. S2C). By contrast, sugar- and fat-conditioned cohorts exhibited overlapping trajectories in feed intake and body composition (Fig. S2E-H). Despite comparable caloric intakes during the intervention period between the SFA- and PUFA-conditioned groups (Fig. S2I), SFA-conditioned mice experienced greater feed efficiency than their counterparts (Fig. S2J), as evidenced by steeper body weight gain relative to feed intake (Fig. S2K) and higher fat mass (Fig. S2L). This is consistent with a broad range of experimental and epidemiological data suggesting increased effective caloric gains under SFA versus PUFA ingestion (*27*).

### Dietary conditioning shapes host brain metabolite profiles

Building on weight-based and time-based measures of eating behavior, we hypothesized that dietary conditioning would differentially impact neuromodulatory pathways in the brain. To test this, we performed targeted and untargeted LC-MS-based metabolomics to identify metabolic responses to diet. Our targeted panel of metabolites spanned monoaminergic and amino acid-derived neurotransmitters, endocannabinoid lipid mediators, and bile acid- and cholesterol-derived signaling molecules (Table S5), and we complemented this targeted analysis with untargeted screening to identify broader metabolic responses to diet.

Targeted metabolomics revealed that dietary fat drives distinct alterations in brain metabolites. In the fat vs. sugar comparison, fat conditioning elevated cholesterol and the endocannabinoids 2-arachidonoylglycerol (2-AG) and anandamide (AEA) (Fig. 2A), whereas sugar conditioning increased GABA (Fig. 2A). In the SFA versus PUFA contrast, PUFA-conditioned mice exhibited elevated levels of the primary serotonin catabolite 5-HIAA, whereas SFA-conditioned mice exhibited elevated cholic acid, pointing to divergent modulation of monoaminergic and bile acid signaling (Fig. 2B).

**Fig. 2:**
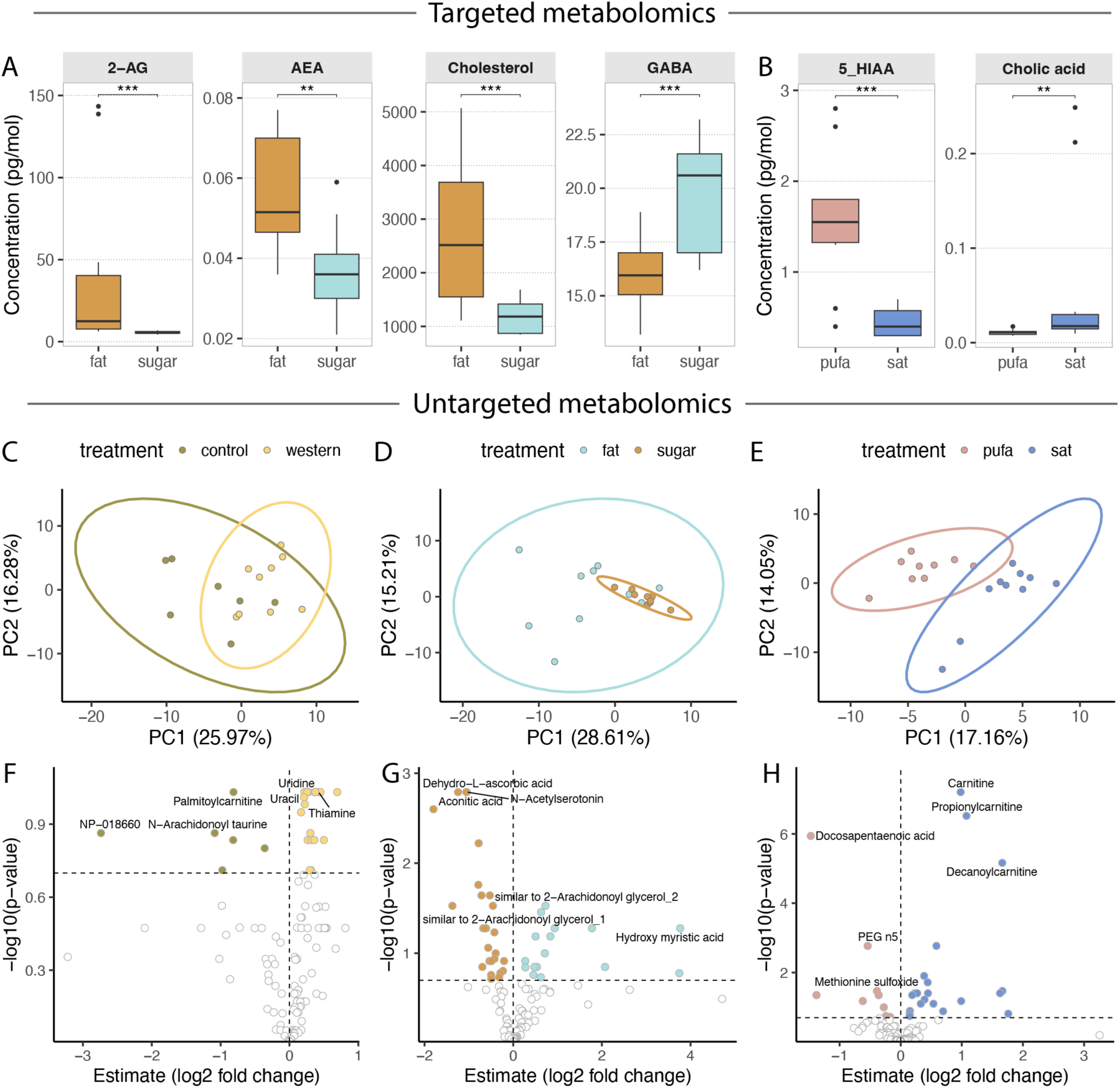
Dietary conditioning induces distinct brain metabolite profiles. **A-B** Targeted brain metabolites that were differentially abundant after dietary conditioning. High-fat diet conditioning elevated brain levels of neuroactive endocannabinoids (2-AG, AEA) and cholesterol, while high-sugar diet conditioning increased GABA (**A**). PUFA conditioning elevated the primary serotonin catabolite 5-HIAA, while SFA conditioning increased cholic acid (**B**). Wilcoxon rank-sum test; *\*\*\* adj-p <* 0.05*, ** adj-p <* 0.1, N.S. = not significant. **C-E** Principal component analysis (PCA) of brain metabolomics data differentiated treatment groups in the Western vs. control (**C**), fat vs. sugar (**D**), and SFA vs. PUFA (**E**) dietary contrasts. Ellipses represent 95% confidence intervals for each group based on the first two principal components. **F-H** Volcano plots showing the distribution of untargeted brain metabolites based on log_2_ fold-change and -log_10_(adj-p) for the Western vs. control (**F**), fat vs. sugar (**G**), and SFA vs. PUFA (**H**) dietary contrasts. Colored data points represent metabolites that differed in abundance between treatment groups; horizontal and vertical dashed lines indicate the significance (*adj-p <* 0.2) and fold-change (log_2_FC > 0) thresholds, respectively. Labeled points denote the top three differentially abundant untargeted metabolites in each contrast.

To illuminate other non-targeted brain metabolites that were affected by dietary conditioning, we complemented our targeted analyses with untargeted metabolomics. In the Western versus control contrast, dietary conditioning led to shifts in brain metabolites involved in energy metabolism, amino acid turnover, neurotransmission, and xenobiotic processing, reflecting the broad neurochemical impact of a fat- and sugar-rich, energy-dense diet (Fig. 2C, 2F; Table S5). In the fat vs. sugar contrast, fat conditioning led to suppression of amino acid and nucleotide metabolites, along with increased lipid-derived signaling molecules and redox compounds (Fig. 2D, 2G; Table S5). In the most granular contrast, SFA conditioning increased levels of medium chain acylcarnitine and fatty acid transport metabolites, alongside changes in neuroactive compounds (Fig. 2E, 2H; Table S5).

### Brain metabolites mediate effects of dietary conditioning on host appetite

We next used mediation analysis (*28*) to test whether brain metabolites may mediate the effects of dietary conditioning on appetite. Mediation analysis quantifies the extent to which the relationship between two variables of interest is influenced by a third variable (mediator) that functions as an intermediary. Here, *mediation effect* refers to the effect of dietary conditioning on appetite that is mediated by brain metabolites, whereas *total effect* refers to the overall effect of dietary conditioning on appetite, regardless of whether it is mediated by brain metabolites (Fig. 3A).

**Fig. 3:**
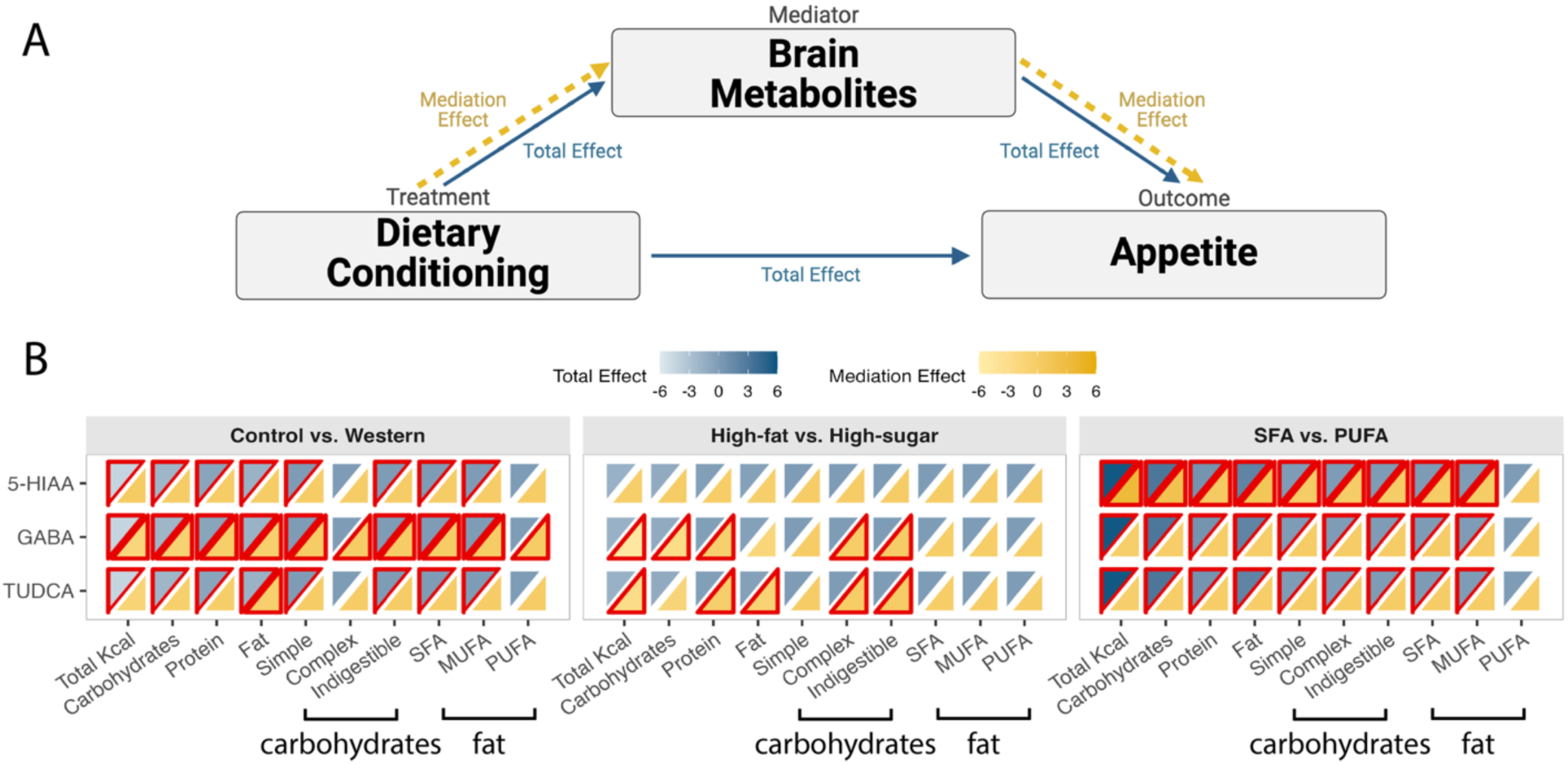
Brain metabolites mediate the effects of dietary conditioning on host appetite. **A** Conceptual framework for the mediation analysis. *Mediation effect* refers to the effect of dietary conditioning on appetite that is mediated by brain metabolites; *total effect* refers to the overall effect of dietary conditioning on appetite, regardless of whether it is mediated by metabolites. **B** Heatmap depicts both the total effect (blue scale) and the mediation effect (yellow scale) of each metabolite on appetite across the Western vs. control, fat vs. sugar, and SFA vs. PUFA dietary contrasts. In the Western vs. control contrast, GABA mediated effects on appetite across all intake levels. In the fat vs. sugar contrast, GABA and TUDCA mediated effects on total caloric intake, and the intakes of protein, fat, and complex and indigestible carbohydrates. In the SFA vs. PUFA contrast, 5-HIAA mediated effects on total caloric intake and the intakes of SFA and MUFA. 5-HIAA also mediated macronutrient- and carbohydrate-specific intake; however, these outcome variables scale proportionally with total caloric intake because these diets were designed to be matched for macronutrients, so their effect sizes are also proportional to total caloric intake. Color scales reflect the magnitudes and directions of total effect and mediation effect; significant effects (*adj-p <* 0.2) are outlined in red.

The total effects of dietary conditioning on appetite were most pronounced in the Western vs. control and SFA vs. PUFA contrasts, whereas the fat vs. sugar contrast showed minimal effects, mirroring the patterns observed in the appetite analyses (Fig. 1). Modeling diet–metabolite and metabolite–appetite relationships separately to examine mediation effects identified 5-HIAA, GABA, and the primary bile acid tauroursodeoxycholic acid (TUDCA) as the brain metabolites that most strongly mediated the effects of dietary conditioning on host appetite whether indexed as general appetite, macronutrient-specific appetite, or substrate-specific appetite (Fig. 3B, Fig. S3). The effects of 5-HIAA, GABA, and TUDCA all showed positive estimates (where control, high-sugar, and SFA diets served as reference groups and Western, high-fat, and PUFA diets served as treatment groups), indicating that these brain metabolites act to increase appetite.

In addition to 5-HIAA, GABA, and TUDCA, we observed a suite of other neuroactive metabolites – including endocannabinoids, biogenic amines, and steroid derivatives – that exerted effects on appetite (Fig. S3). However, the effects of these additional metabolites were less consistent and more context-specific, so we elected to focus downstream work on 5-HIAA, GABA, and TUDCA as the most promising appetite-modulatory metabolites.

### Diet remodels gut microbial composition and function linked with 5-HIAA

5-HIAA (*29, 30*), GABA (*31–33*), and TUDCA (*34*) are all known to interact with the gut microbiota. Microbes promote 5-HIAA production during early-life nutrition (*29*) and enable stress-induced serotonergic responses (*30*). Specific microbial strains also synthesize (*32, 33*) and consume (*31*) GABA, shaping neurochemical signaling through cross-feeding interactions. Meanwhile, gut dysbiosis reduces TUDCA biosynthesis, weakening its anti-inflammatory and metabolic regulatory roles (*34*). Given these effects, and since diet is known to be a principal determinant of the gut microbiome (*35, 36*), we examined how dietary conditioning altered gut microbial composition.

Bray-Curtis principal component analysis revealed structural differences in gut microbial communities across all three dietary contrasts, confirming that dietary exposure robustly shapes microbial ecology (Fig. 4A-C). To characterize these shifts, we conducted ANCOM-BC2-based differential abundance analysis (*37*) and identified distinct microbial signatures associated with each diet. Western diet conditioning enriched diverse taxa across the orders *Clostridiales*, *Erysipelotrichales*, *Bacteroidales*, and *Lactobacillales* (Fig. 4D), while high-sugar conditioning selectively enriched the family *Christensenellaceae* (Fig. 4E). SFA and PUFA diets induced distinct microbial shifts within the order *Clostridiales*, with SFA conditioning favoring several *Ruminococcus* and *Peptostreptococcaceae* lineages, and PUFA conditioning enriching members of the *Lachnospiraceae* family (Fig. 4F), a group that includes known Stickland fermenters capable of oxidizing tryptophan into neuroactive metabolites (*38*).

**Fig. 4:**
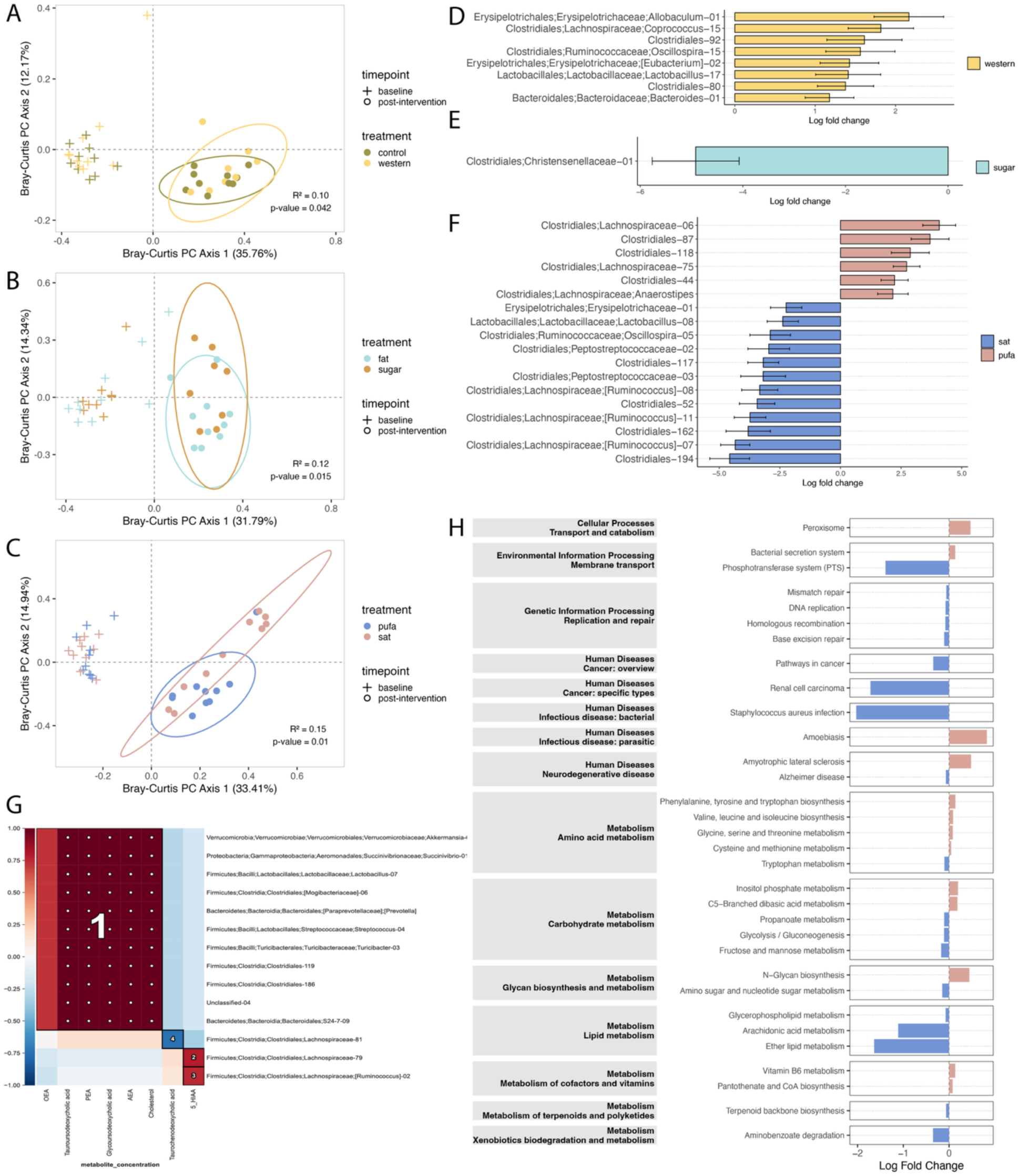
Dietary conditioning remodels gut microbiome composition and functions. **A-C** Bray-Curtis principal coordinate plots illustrate diet-induced shifts in gut microbial community structure. Significant separation of post-intervention gut microbiota community structure was observed in all diet contrasts: Western vs. control (**A**), fat vs. sugar (**B**), and SFA vs. PUFA (**C**). PERMANOVA; *p*-values shown in panel. **D-F** Differentially abundant microbial taxa after dietary conditioning as indexed by ANCOM-BC2. Enriched taxa in the Western-conditioned group included Firmicutes (e.g., *Lachnospiraceae*, *Ruminococcaceae*, *Erysipelotrichaceae*, *Lactobacillaceae*) and Bacteroidetes (*Bacteroidaceae*). (**D**). High-sugar diet conditioning enriched the *Christensenellaceae* family (**E**). SFA and PUFA diets induced distinct microbial shifts within the order *Clostridiales*, with PUFA conditioning enriching several members of the *Lachnospiraceae* family and SFA conditioning favoring the genus *Ruminococcus* (**F**). **G** HAllA revealed distinct clusters of metabolite–microbe associations, with bile acids broadly associating with diverse taxa (Block 1) and 5-HIAA selectively associating with *Lachnospiraceae* (Blocks 2– 3). **H** PICRUSt2-based metagenomic inference identified diet-specific enrichment of KEGG pathways, with SFA conditioning enriching pathways involved in glycan metabolism and infectious disease, and PUFA conditioning increasing pathways related to neurodegenerative disease and amino acid metabolism, including tryptophan biosynthesis.

To test whether these diet-induced microbial changes might relate to brain signaling, we used HAllA (*39*) and identified robust associations between brain metabolite levels and gut microbial taxa (Fig. 4G). Strikingly, the metabolites linked to microbial composition were largely distinct from those identified as mediators of appetite (Fig. 3, Fig. S3). While bile acids dominated the HAllA-associated set, none appeared as mediators, suggesting their role as microbially derived signaling molecules rather than modulators of feeding behavior. In contrast, 5-HIAA emerged as a point of overlap, such that elevated 5-HIAA levels in PUFA-conditioned mice corresponded with both changes in gut microbiota composition (increased *Lachnospiraceae*) and increased appetite relative to SFA-conditioned mice.

To understand whether elevated 5-HIAA levels might arise due to changes in the functional potential of the gut microbiome, we employed PICRUSt2-based metagenome inference (*40*). Whereas SFA-conditioned mice showed enrichment in categories related to glycan biosynthesis and metabolism and infectious disease, we found that PUFA-conditioned mice exhibited enrichment in KEGG pathway categories related to neurodegenerative diseases and amino acid metabolism, including enrichment of tryptophan metabolic pathways (Fig. 4H). This suggests that PUFA conditioning increased microbial capacity to supply tryptophan, a key substrate for host serotonin biosynthesis (*41*).

Taken together, these data position 5-HIAA as a unique molecular node linking microbial composition, brain signaling, and increased appetite after PUFA conditioning.

### Gut microbes causally influence host appetite

Building on our discovery that 5-HIAA links gut microbial composition and function to appetite regulation, we next asked whether the gut microbiome itself plays a causal role in shaping host appetite. To test this, we transplanted into germ-free C57BL/6J mice pre-PUFA-conditioning (PUFA_0_) and post-PUFA-conditioned (PUFA_+_) gut microbiota harvested from murine donors in the SFA vs. PUFA experiment (Fig. 5A). Inoculated mice were subjected to baseline and post-inoculation 24-hour intake assays, with appetite indexed by total, macronutrient-specific, and substrate-specific intake, mirroring the study design used in conventional mice.

**Fig. 5:**
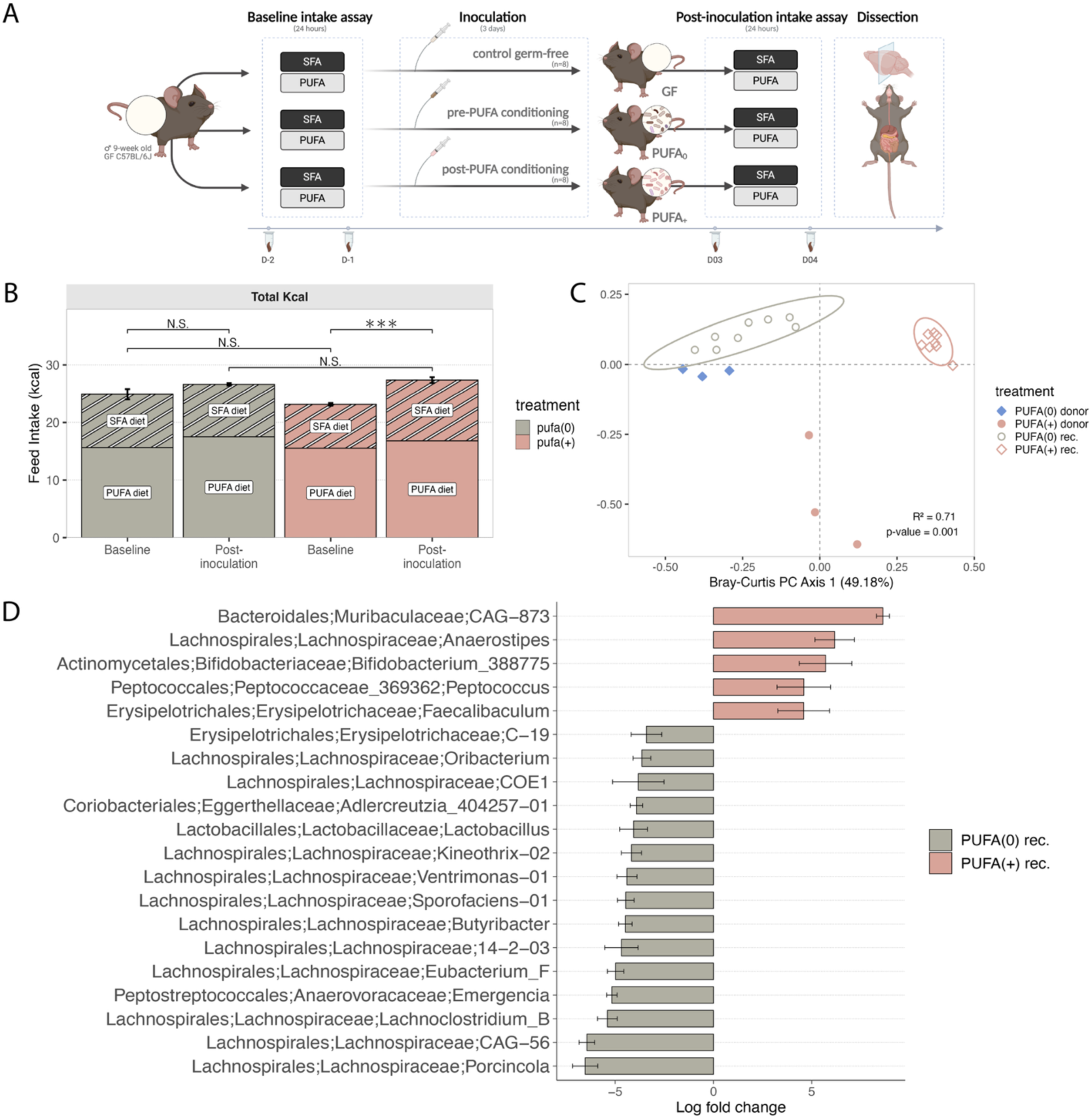
PUFA-conditioned gut microbiome carries an appetite-promoting signature. **A** Schematic of the gnotobiotic mouse experiment. Image created in BioRender. **B** Caloric intake during the 24-hour intake assays conducted at baseline and post-inoculation (n = 8/group). Inoculation led to higher total caloric intake in PUFA_+_ but not PUFA_0_ recipients. Wilcoxon rank sum test, *\*\*\* adj-p <* 0.05*, ** adj-p <* 0.1*, * adj-p <* 0.2, N.S. = not significant. **C** Bray-Curtis principal coordinate plot illustrates transmission of donor gut microbiota into recipients and confirms that PUFA_0_ and PUFA_+_ recipients harbored distinct gut microbial communities. PERMANOVA, *p <* 0.05. **H** ANCOM-BC2-based differential abundance analysis identified taxa enriched in PUFA_+_ versus PUFA_0_ recipients, including several taxa from the family *Lachnospiraceae*.

Consistent with PUFA-conditioned microbiota promoting increased appetite, PUFA_+_ recipients exhibited higher total caloric intake than PUFA_0_ recipients after inoculation (Fig. 5B), echoing the enhanced appetite observed in PUFA-conditioned conventional mice post-intervention (Fig. 1E). Beta-diversity analysis confirmed successful engraftment of the donor microbiome into their respective recipients and illustrated distinct gut microbiota profiles between PUFA_0_ and PUFA_+_ recipients (Fig. 5C), with the family *Lachnospiraceae* among the taxa enriched in both the PUFA-conditioned donors (Fig. 4F) and PUFA_+_ recipients (Fig. 5D). These data highlight the gut microbiome’s causal influence in modulating host appetite and suggest an appetite-promoting microbial signature dominated by *Lachnospiraceae*.

## DISCUSSION

In our murine model, PUFA conditioning promoted increased caloric intake and elevated brain 5-HIAA, a serotonin catabolite that was found to mediate the effects of dietary conditioning on appetite. Levels of 5-HIAA in the brain were associated with *Lachnospiraceae* abundance, and PUFA conditioning enriched predicted microbial pathways related to serotonergic signaling, prompting us to examine the microbiome as a potential mechanism. Transplantation of a PUFA-conditioned microbiome (PUFA_+_) was sufficient to promote increased caloric intake in murine gnotobiotic recipients, indicating a transmissible microbiome-dependent behavioral phenotype.

PUFA conditioning robustly enhanced appetite, as indexed by total caloric intake in 24-hour intake assay conducted post-intervention. Concordantly, our behavioral analysis revealed that PUFA-conditioned mice consumed more pellets while spending comparable time at the feeder, indicating a higher eating rate – a pattern known to increase ad libitum energy intake (*42*). While rodent studies, including ours, have noted a trend toward greater energy intake in response to PUFA feeding compared to SFA feeding (*43–46*), human studies report more variable outcomes: some find that PUFA-rich meals suppress appetite more effectively than SFA or MUFA meals (*47–49*), while others observe no effect of fatty acid saturation or chain length on energy intake (*50, 51*). These mixed findings may reflect the consumption by humans of mixed meals versus purified diets, reliance on subjective appetite ratings versus direct measurements of caloric intake, and the short-term nature of many human feeding trials relative to the conditioning paradigms used in animal models.

Consumption of a PUFA-rich diet elevated brain levels of 5-HIAA, the primary serotonin catabolite (*52*), which was subsequently found to mediate the effect of dietary conditioning on host appetite. Classic models attribute serotonin-driven appetite regulation to carbohydrate-induced shifts in plasma tryptophan availability (*53–55*), the precursor to serotonin. Although we observed diet-induced shifts in the gut microbiome that enriched for tryptophan biosynthetic pathways, we found no differences in brain tryptophan levels between dietary groups. Instead, the elevation of 5-HIAA following PUFA conditioning suggests increased serotonin turnover as a downstream mechanism. Because peripheral serotonin does not cross the blood-brain barrier (*56*), we propose that PUFA ingestion stimulates central serotonin turnover through indirect mechanisms such as gut-derived vagal signaling. This is supported by the enrichment of *Lachnospiraceae* – members of which conduct Stickland fermentations, including oxidative metabolism of tryptophan into neuroactive catabolites (*38*) – in microbiota profiles known to engage vagal monoaminergic circuit (*57*). While prior reports have linked PUFA consumption to increased serotonergic turnover (*58–60*), our study is unique in proposing microbiome-mediated gut-brain signaling as a pathway through which dietary fats shape serotonergic tone.

Our experiments in gnotobiotic mice suggest that PUFA-mediated shifts in the gut microbiome were sufficient to drive appetite changes in the absence of direct dietary conditioning. Recapitulation of behavioral effects in microbiota transplant recipients adds to growing evidence that the gut microbiome exerts causal influence over host appetite and nutrient preference (*15, 61–63*), and that diet-derived microbial metabolites can regulate host appetite via gut-brain signaling (*64*). By demonstrating a new PUFA-serotonergic pathway mediating microbiome-dependent appetite enhancement, we provide biological validation of food-microbe-metabolite interactions previously only predicted using deep-learning approaches (*65*). Furthermore, our integration of brain metabolomics, microbiome profiling, and gnotobiotic experimentation delivers mechanistic granularity and translational insights, complementing clinical studies that link dietary modulation of gut microbiota to host energy balance (*66*). Together, these findings position the gut microbiome as both a target and transmitter of dietary and neural cues in the regulation of appetite.

While our findings outline a diet-microbiome-gut-brain axis shaping PUFA-driven modulation of appetite, further work to enhance mechanistic resolution will be valuable. For instance, while 16S rRNA gene sequencing was appropriate to establish the dietary sensitivity of the gut microbiota, validate the fidelity of fecal microbiota transplantation, and enable PICRUSt2-based prediction of metagenomic pathway enrichment, shotgun metagenomic sequencing will be necessary to assay strain-level variation and changes in functional capacity within PUFA-conditioned microbial communities. The use of synthetic microbial communities could facilitate mechanistic dissection of whether and how *Lachnospiraceae* increases tryptophan availability to shape host eating behavior. Additionally, while our data positions 5-HIAA as a key metabolite linking PUFA consumption, gut microbiome profile, and feeding behavior, functional validation remains essential. Future studies leveraging neural tools – e.g., calcium imaging, *Fos* expression mapping, and targeted gain/loss-of-function approaches – will be critical to confirm whether serotonergic circuits mediate the behavioral effects of a PUFA-conditioned gut microbiome. Collectively, these future directions will help define how consumption of dietary fats shapes appetite in industrialized populations experiencing alterations in dietary composition and elevated rates of both disordered eating and metabolic disease.

## MATERIALS AND METHODS

### Animals and experimental design

Eight-week-old male C57BL/6J mice were used in all conventional and gnotobiotic experiments. Conventional mouse studies were conducted at the Biological Research Infrastructure (BRI) research facility at Harvard University under Harvard University Institutional Animal Care and Use Committee (IACUC) protocol #17-06-306. The gnotobiotic mouse experiment was conducted at the Massachusetts Host-Microbiome Center at Brigham and Women’s Hospital under Harvard Medical School IACUC protocol #2016N000141. Mice were housed individually at_25°C on a 12h:12h light:dark cycle and had ad libitum access to water and food, as specified below.

### Food intake assay and dietary conditioning

To assess how dietary exposures influence appetite and brain-gut interactions, we implemented a baseline intake assay, followed by a two-week dietary conditioning protocol, followed by a post-conditioning intake assay.

During each intake assay, mice were provided with simultaneous access to two diets for 24 hours, enabling quantification of both total food intake and preferences within a dietary contrast. Each dietary pairing represented one of three contrasts relevant to an industrialized diet: Western vs. control (TD.110919 vs. TD.140806, **Table S2**), high-fat vs. high-sugar (TD.220629 vs. TD.220628; **Table S3**), and SFA-rich vs. PUFA-rich (TD.220630 vs. TD.220631; 34.5% kcal from fat for both; **Table S4**). Video recordings taken during the intake assays using overhead Raspberry Pi cameras were used to validate food intake measurements. During the two-week dietary conditioning period, animals received one of the two diets from their assigned dietary contrast, with body weight and feed intake recorded every other day.

We quantified appetite at three levels: (i) general appetite, indexed by overall caloric consumption; (ii) macronutrient-specific appetite, indexed by consumption of calories derived from protein, carbohydrate, or fat; and (iii) substrate-specific appetite, indexed by consumption of calories derived from key classes of carbohydrate (simple carbohydrate, complex carbohydrate, or indigestible fiber) or fat (SFA, MUFA, or PUFA). In the Western vs. control and fat vs. sugar contrasts, the diets differ in nutrient composition, allowing meaningful deconstruction of total intake at all three levels. In comparison, the two diets in the third contrast (SFA vs. PUFA) are isocaloric and matched in their macronutrient composition, differing only in fatty-acid profiles. Because the diets are matched for all nutrients except fat type, macronutrient- and carbohydrate-specific intake values are proportional to total intake by design, precluding meaningful analyses of macronutrient-specific and carbohydrate-specific preferences. We have therefore focused on general appetite and fatty acid-specific appetite for this contrast.

### Behavioral tracking

During the 24-hour intake assay, the behaviors of the experimental mice were recorded using a Raspberry Pi camera installed at the top of the cage. The camera was programmed to record for 24 hours, and the videos were analyzed using DeepLabCut, a deep learning-based software for markerless pose estimation based on a deep neural network. To label and predict the coordinates of each body part with DeepLabCut, we extracted 240 characteristic frames and labeled the nose point and tail point. The extracted images were transformed into training sets on which the deep neural network was trained. The network comprised ResNet-50 and deconvolutional layers, the outputs for which are score maps representing the soft predictions for the location of each body part. The network was trained through 800,000 iterations, minimizing the cross-entropy of the predicted probability distribution relative to the ground-truth probability distribution. The trained model was evaluated by computing the mean average Euclidean error between the manual labels and the labels predicted by DeepLabCut. The trained model achieved a 97.18% accuracy. After training the model, the body part coordinates were predicted. The coordinates were used to calculate the time spent in each zone. Consummatory behavior was defined as nose point touching the feeder for more than 2 seconds but less than 60 seconds. All other behaviors (resting, grooming, etc.) were classified as less than 60 seconds. Drinking was defined as nose point touching the water valve for 0.1 seconds or more. The cumulative time spent between the two feeders was used as a proxy for intake, complementing the endpoint feed intake measurement.

### Metabolomics

To identify brain metabolic signatures associated with dietary conditioning, we performed targeted and untargeted metabolomics analyses on homogenized brain tissue using LC-MS. Brain samples from the conventional mouse experiments (n = 57) were collected and stored at –80 °C. One hemisphere was homogenized in acetonitrile:methanol (1:1, v/v; LC-MS grade) using a bead beater (TissueLyser LT, 10 min, 50 Hz). Two aliquots were collected for metabolite extraction: one for targeted neurotransmitter analysis (NeuroT method) and one for targeted non-polar metabolites and untargeted profiling (PFPP method). Targeted compounds were selected to encompass key neuroactive metabolites and signaling lipids across monoaminergic, glutamatergic, endocannabinoid, and bile acid pathways implicated in nutrient sensing and appetite regulation.

Internal standards were added before extraction. Samples were vortexed (1 min), sonicated (10 min), and centrifuged (18,000 rcf, 15 min), with supernatants transferred to low-binding tubes. NeuroT samples were exclusively used for targeted analysis and underwent dansyl chloride derivatization (4 mg/mL, 30 min, 37 °C) and were quenched with 30% formic acid. These samples were analyzed using an AB Sciex 7500 triple quadrupole mass spectrometer coupled with an Agilent 1290 LC, employing multiple reaction monitoring (MRM) on a Kinetex Polar C18 column (150 × 2.1 mm, 2.6 µm) at 45 °C. The mobile phase consisted of 0.1% formic acid in water (A) and in acetonitrile (B), with a gradient elution from 5% to 100% B over 8 min, held until 12 min. PFPP samples were used for both targeted quantification and untargeted profiling. Samples were dried under nitrogen, resuspended in methanol:acetonitrile (1:1, v/v), vortexed, and centrifuged. Targeted analytes were quantified using an in-house standard curve and batch-corrected linear regression. For untargeted analysis, 5 µL injections were analyzed on a ThermoFisher ID-X Tribrid mass spectrometer coupled to a Vanquish LC. Chromatographic separation was achieved on a Restek Allure PFPP column (150 × 2.1 mm, 5 µm, 35 °C) using a 34-minute gradient of acetonitrile:isopropanol (9:1, v/v). AquireX deep MS2 scanning was applied to pooled QC samples. Untargeted data were processed using Compound Discoverer 3.3. Peak extraction, alignment, gap-filling, and background subtraction were followed by normalization based on resuspension volume and median centering. Compound identities were assigned through mzCloud and in-house libraries, with manual curation. Metabolites were subsequently classified based on their biological roles with reference to the Chemical Entities of Biological Interest (ChEBI) database. Statistical analyses included t-tests with Benjamini-Hochberg correction (FDR < 0.01) and partial least squares discriminant analysis (PLS-DA).

### 16S rRNA gene amplicon sequencing

To characterize gut microbial composition, we performed 16S rRNA gene amplicon sequencing on fecal samples. Fecal DNA was extracted using the E.Z.N.A. Soil DNA Kit (Omega Bio-Tek) and PCR-amplified with custom-barcoded 515F and 806R primers targeting the V4 region of the 16S rRNA gene. PCR was performed on each sample in triplicate, including sample-specific negative controls, on a BioRad T100 thermocycler using the following protocol: 95°C for 3 min; 35 cycles of 94°C for 45 s, 50°C for 30 s, and 72°C for 90 s; followed by a final extension at 72°C for 10 min. Triplicate reactions per sample were pooled, and amplification was confirmed by 1.5% agarose gel electrophoresis. Amplicons were purified using Ampure XP beads, quantified with the Quant-iT PicoGreen dsDNA Assay Kit (Invitrogen), and pooled in equimolar amounts. Sequencing (2 x 150 bp) was performed on an Illumina NovaSeq X Plus, generating 171,830 ± 3,740 SE (conventional study) and 104,444 ± 12,034 (gnotobiotic study) reads per sample. Raw FASTQ files were processed in QIIME2 (*67*) (version 2023.7) using the DADA2 plugin, with reads truncated at 150 bp to optimize sequence quality. Taxonomic classification of amplicon sequence variants (ASVs) was performed using the Greengenes2 (*68*) database, and a rooted phylogenetic tree was constructed to support downstream diversity analyses. The ASV feature table, taxonomy, and phylogeny were then imported into R (version 4.3.2) using the qiime2R package (version 0.99.6). Subsequent data processing and diversity analyses were conducted using the phyloseq (*69*) package (version 1.46.0). Differential abundance testing was performed using ANCOM-BC2 (*37*), and microbiota–metabolite associations were evaluated using HAllA (*39*); see Statistical Analysis section for further details. PICRUSt2 (*40*) was used to predict functional pathways, and the output was analyzed with MaAsLin2 to compare gene pathways between treatment groups using the ggpicrust2 package (*70*) in R (version 1.7.3).

### Gnotobiotic experiment

To evaluate the causal contribution of the gut microbiome to diet-induced changes in appetite, we conducted a gnotobiotic experiment in which germ-free mice were inoculated with fecal microbiota from conventional mice before and after their exposure to PUFA diet conditioning. To generate pre-PUFA exposure and post-PUFA exposure inocula, we pooled fecal samples collected from the same conventional donor mice (n=3) before and after two weeks of PUFA-based dietary conditioning. Pooled feces were diluted 1:30 in reduced PBS, vortexed and spun down, with the supernatant used as inocula. To generate sterile control inocula, we autoclaved aliquots of these same preparations. 24 germ-free C57BL/6J mice (8–10 weeks old) were divided evenly into three groups and gavaged with 393.9 μl of the pre-PUFA-conditioned inocula (PUFA_0_), 195.5 μl post-PUFA-conditioned inocula (PUFA_+_), and 144.9 μl sterile control inocula (control). The sample size for our gnotobiotic study was determined based on *a priori* power analyses using the pwr package in R (v1.3.0), indicating that a sample size of 8 recipient mice per group would give us 90% power to detect an effect size = 0.8 at α = 0.05.

All recipient mice underwent a baseline 24-hour intake assay, with simultaneous access to isocaloric SFA (34.5% kcal from fat, AMF) and PUFA (34.5% kcal from fat, flaxseed oil) diets (Table S4). Mice were then colonized via gavage and allowed three days for the microbiota to stabilize, after which the intake assay was repeated. Diet intake was recorded at the start and end of each 24-hour assay to quantify appetite. Appetite was evaluated by summing the intake of both diets to determine total caloric intake. Diets were then deconstructed into macronutrient- and substrate-specific components for higher-resolution analysis.

### Statistics

Wilcoxon rank-sum tests were used to compare appetite across groups and time points. We used the mediation package (*28*) to test whether brain metabolite profiles mediated the effect of dietary conditioning on host appetite. This pipeline implements a nonparametric bootstrap to estimate average causal mediation effects, enabling inference by decomposing total effects into direct and metabolite-mediated components, with statistical significance defined at adjusted p-value < 0.2. Differential abundance testing was performed using the ANCOMBC package (v2.0.3) (*37*), implementing Analysis of Compositions of Microbiomes with Bias Correction 2 (ANCOM-BC2), a method that adjusts for sampling fraction bias and provides valid inference with adjusted p-values and confidence intervals. To identify structured associations between gut microbial taxa and metabolite profiles, HAllA (Hierarchical All-against-All association testing) (*39*) was applied to detect correlated features while controlling for multiple hypothesis testing, using q-value < 0.05 to define significance. Adjusted p-values were calculated using the Benjamini–Hochberg method. All statistical analyses were conducted using R version 4.2.2.

## LIST OF SUPPLEMENTARY MATERIALS

Fig. S1: Intake of macronutrients and nutrient subclasses during the post-intervention 24-hour intake assay.

Fig. S2: Body composition and feed intake trajectory during the diet conditioning period.

Fig. S3: Full spectrum of brain metabolite contributions to appetite regulation.

Table S1: Macronutrient composition of experimental diets.

Table S2: Nutritional composition of Western and control diets.

Table S3: Nutritional composition of high-fat and high-sugar diets.

Table S4: Nutritional composition of SFA and PUFA diets.

Table S5: Differentially abundant untargeted metabolites and their classes.

## ACKNOWLEDGEMENTS

We thank the members of the Carmody Lab for their feedback on early drafts of this manuscript, and Charles Vidoudez for his assistance with metabolomics. This work was supported by grants from the Japan Society for the Promotion of Science 23KJ0457 (to YJL), P30 DK034854 and R01 AI179807 (to LB), and the William F. Milton Fund and Harvard Dean’s Competitive Fund for Promising Scholarship (to RNC).

## AUTHOR CONTRIBUTIONS

Conceptualization: YJL, RNC. Methodology: YJL, RNC, MT. Investigation: YJL, YV. Statistical analysis: SE, YJL. Visualization: SE, YJL. Funding acquisition: YJL, RNC. Project administration: YJL. Supervision: RNC. Writing – original draft: YJL. Writing – review & editing: YJL, SE, MT, LB, RNC.

## DATA AND MATERIALS AVAILABILITY

16S rRNA sequencing data generated in this study have been deposited in the NCBI Sequence Read Archive under BioProject accession PRJNA1291436. Targeted and untargeted metabolomics data have been deposited to the Metabolomics Workbench under DataTrack ID 6193 and are currently under curation. Additional data and analysis code are available upon request.

