## Supplementary Materials for "Polyunsaturated fatty acids promote appetite via the microbiome-gut-brain axis"

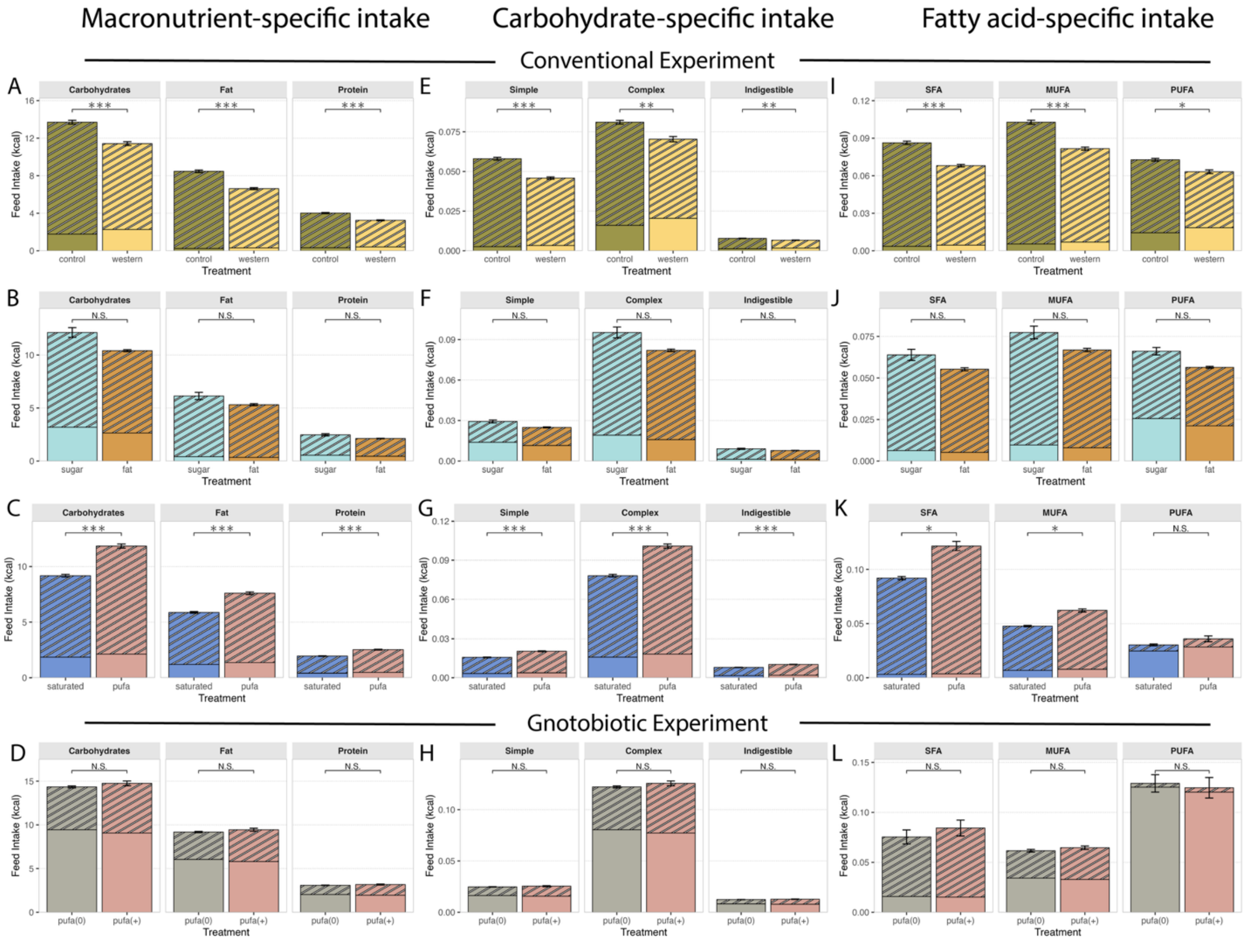


**Fig. S1: Intake of macronutrients and nutrient subclasses during the post-intervention 24-hour intake assay. A-D** Macronutrient-specific intake: Western and SFA diet conditioning suppressed intake of carbohydrates, fat, and protein. **E-H** Carbohydrate-specific intake: Western and SFA diet conditioning suppressed intake across simple, complex, and indigestible carbohydrates. **I-L** Fat-specific intake: Western diet suppressed intake across all fatty acid subclasses; PUFA conditioning selectively increased intake of SFA and MUFA; no observable changes between the PUFA$_0$ and PUFA$_+$ recipients. In panels **C**, **G**, **D**, and **H**, differences between SFA- vs. PUFA-conditioned groups and PUFA$_0$ vs. PUFA$_+$ groups are simply proportional to total caloric intake because these diets were designed to be matched for macronutrients. Wilcoxon rank-sum test; *** *adj-p* < 0.05, ** *adj-p* < 0.1, * *adj-p* < 0.2, N.S. = not significant. Shading denotes dietary source of calories during the assay.

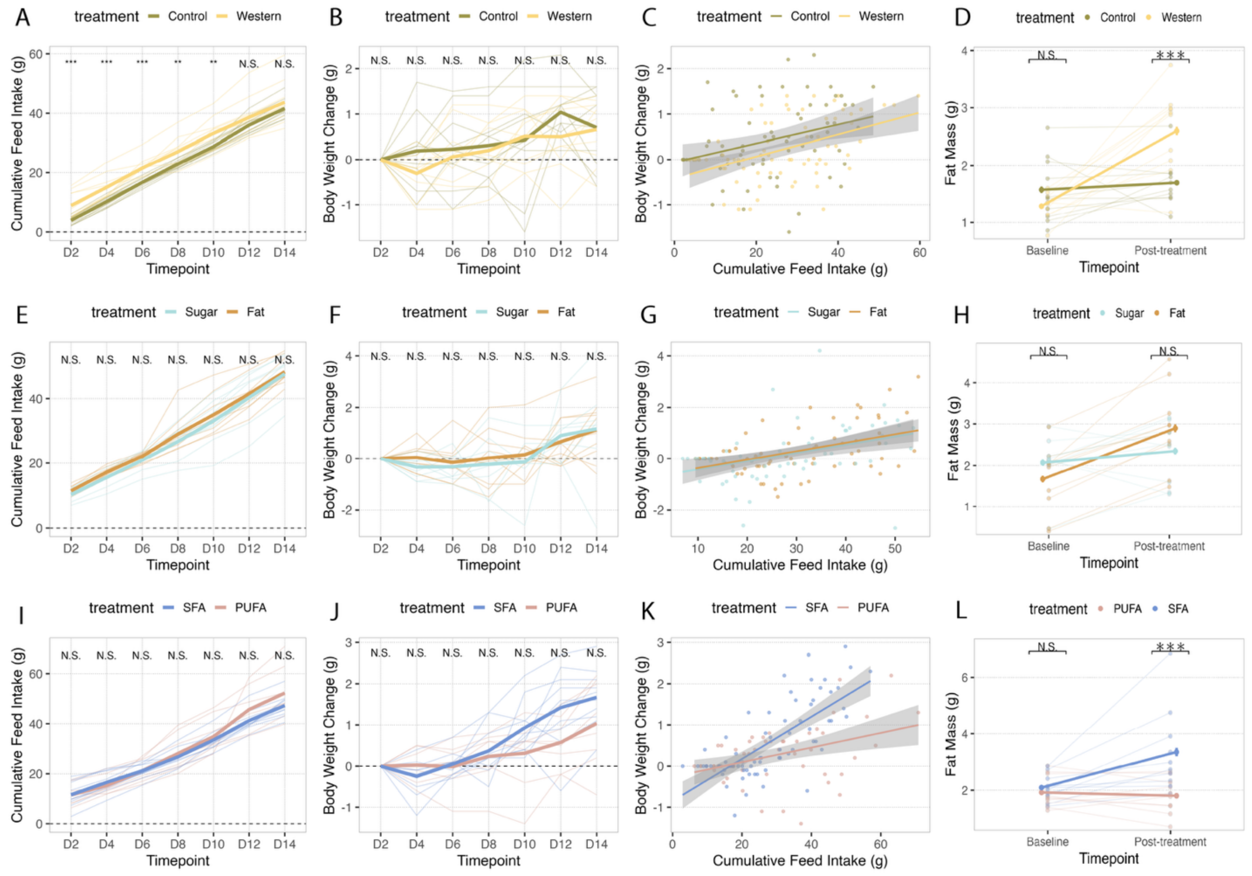


**Fig. S2: Body composition and feed intake trajectory during the diet conditioning period. A, E, I** Cumulative feed intake over the 14-day conditioning period for the control vs. Western (**A**), fat vs. sugar (**E**), and PUFA vs. SFA (**I**) dietary contrasts. **B, F, J** Percent body weight changes relative to day 2 over the 14-day conditioning period across control vs. Western (**B**), fat vs. sugar (**F**), and PUFA vs. SFA (**J**) dietary contrasts. **C, G, K** Linear relationships between cumulative feed intake and body weight change over the 14-day conditioning period across control vs. Western (**C**), fat vs. sugar (**G**), and PUFA vs. SFA (**K**) dietary contrasts. **D, H, L** Body composition before and after the 14-day conditioning period across control vs. Western (**D**), fat vs. sugar (**H**), and PUFA vs. SFA (**L**) dietary contrasts. Western-conditioned (**D**) and SFA-conditioned (**L**) groups gained more fat mass than their counterparts. Wilcoxon rank-sum test; *** *adj-p* < 0.05, ** *adj-p* < 0.1, * *adj-p* < 0.2, N.S. = not significant.

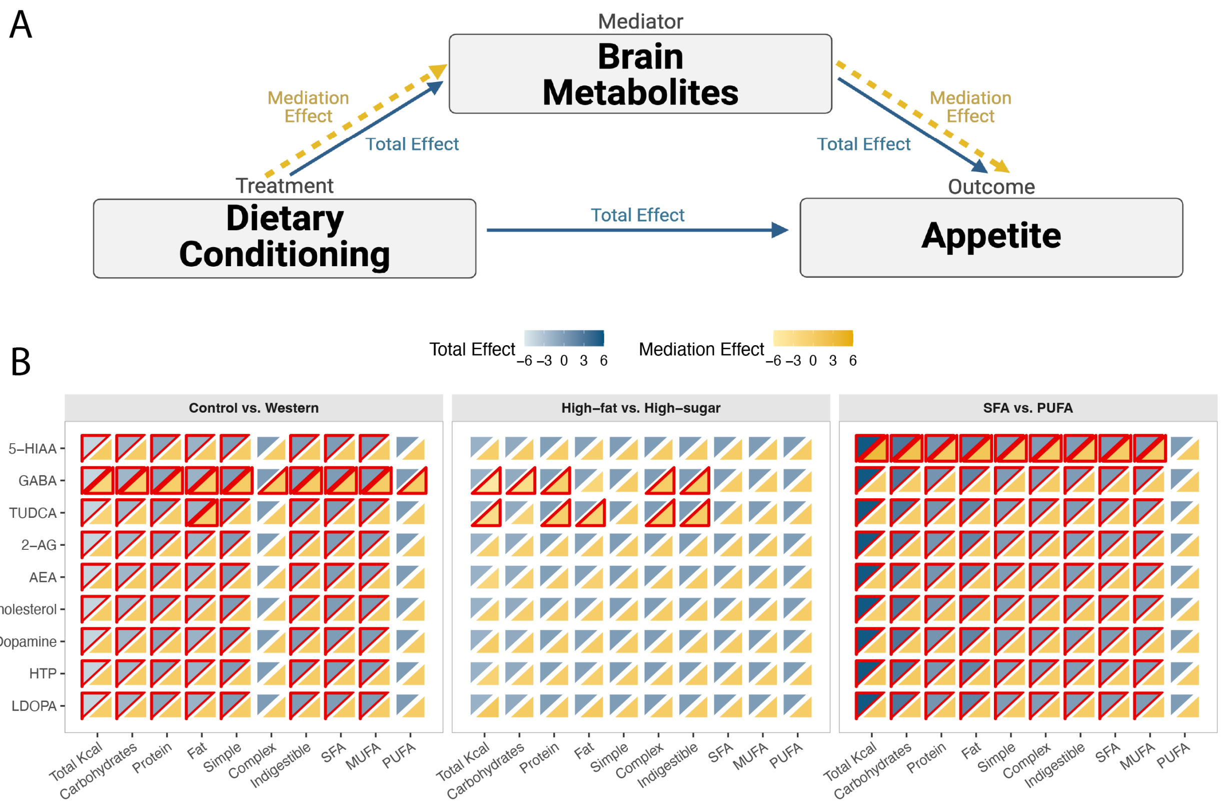


**Fig. S3: Full spectrum of brain metabolite contributions to appetite regulation. A** Conceptual framework for the mediation analysis. *Mediation effect* refers to the effect of dietary conditioning on appetite that is mediated by brain metabolites; *total effect* refers to the overall effect of dietary conditioning on appetite, regardless of whether it is mediated by metabolites. **B** Heatmaps summarize total effects (blue gradient) and mediation effects (yellow gradient) for all quantified brain metabolites across three dietary contrasts (control vs. Western, high-fat vs. high-sugar, SFA vs. PUFA). Metabolites displayed here were those showing overlapping significance in separate tests of diet conditioning→metabolite and metabolite→appetite relationships. This extended panel reveals additional compounds that modulate different dimensions of appetite. Across contrasts, GABA and 5-HIAA emerge as consistent mediators of total caloric intake, while endocannabinoids (2-AG, AEA), biogenic amines (dopamine, HTP, L-DOPA), and sterols (TUDCA, cholesterol) selectively influence total caloric, macronutrient-, and substrate-specific intake. Color intensity reflects effect size and direction for both total and mediation paths; significant effects (*adj-p* < 0.2) are outlined in red.

### Table S1: Macronutrient composition of experimental diets

| | Western TD.110919 | Control TD.140806 | nTWD Fat TD.220629 | Sucrose TD.220628 | SFA TD.220630 | PUFA TD.220631 |
|---|---|---|---|---|---|---|
| **Protein % by weight** | 16.8 | 12.4 | 12.4 | 12.4 | 12.4 | 12.4 |
| **Carbohydrate % by weight** | 54.4 | 68.4 | 57.4 | 71.8 | 57.4 | 57.4 |
| **Fat % by weight** | 16.7 | 4.1 | 16.3 | 4.1 | 16.3 | 16.3 |
| **Protein % kcal from** | 15.5 | 13.7 | 11.6 | 13.2 | 11.6 | 11.6 |
| **Carbohydrate % kcal from** | 50.0 | 75.9 | 53.9 | 76.8 | 53.9 | 53.9 |
| **Fat % kcal from** | 34.5 | 10.3 | 34.5 | 10.0 | 34.5 | 34.5 |
| **kcal/g** | 4.4 | 3.6 | 4.3 | 3.7 | 4.3 | 4.3 |

### Table S2: Nutritional composition of Western and control diets

| Ingredient (g/kg) | Western diet | Control diet |
|---|---|---|
| **Casein** | 190.0 | 140.0 |
| **L-Cystine** | 2.85 | 1.8 |
| **Corn starch** | 230.0 | 460.427 |
| **Maltodextrin** | 70.0 | 155.0 |
| **Sucrose** | 255.604 | 100.0 |
| **Olive oil** | 28.0 | - |
| **Soybean oil** | 31.4 | 40.0 |
| **Corn oil** | 16.5 | - |
| **Lard** | 28.0 | - |
| **Beef tallow** | 24.8 | - |
| **Anhydrous milkfat** | 36.3 | - |
| **Cholesterol** | 0.4 | - |
| **Cellulose** | 30.0 | 50.0 |
| **Mineral mix, nTWD** | 35.0 | - |
| **Sodium chloride** | 4.0 | - |
| **Vitamin mix, nTWD** | 15.0 | - |
| **Choline bitartate** | 2.1 | 2.75 |
| **Vitamin K1, phylloquinone** | 0.003 | 0.002 |
| **Thiamine (81%)** | 0.015 | 0.013 |
| **TBHQ, antioxidant** | 0.028 | 0.008 |
| **Mineral mix, AIN93-M-MX** | - | 35.0 |
| **Vitamin mix, AIN93-VX** | - | 15.0 |
| **Flaxseed oil** | - | - |

### Table S3: Nutritional composition of high-fat and high-sugar diets

| Ingredient (g/kg) | High-fat diet | High-sugar diet |
|---|---|---|
| Casein | 140.0 | 140.0 |
| L-Cystine | 1.8 | 1.8 |
| Corn starch | 338.552 | 355.909 |
| Maltodextrin | 155.0 | 70.0 |
| Sucrose | 100.0 | 309.518 |
| Olive oil | 27.448 | - |
| Soybean oil | 30.782 | 40.0 |
| Corn oil | 16.175 | - |
| Lard | 27.448 | - |
| Beef tallow | 24.312 | - |
| Anhydrous milkfat | 35.685 | - |
| Cholesterol | - | - |
| Cellulose | 50.0 | 30.0 |
| Mineral mix, nTWD | - | - |
| Sodium chloride | - | - |
| Vitamin mix, nTWD | - | - |
| Choline bitartate | 2.75 | 2.75 |
| Vitamin K1, phylloquinone | 0.002 | 0.002 |
| Thiamine (81%) | 0.013 | 0.013 |
| TBHQ, antioxidant | 0.033 | 0.008 |
| Mineral mix, AIN93-M-MX | 35.0 | 35.0 |
| Vitamin mix, AIN93-VX | 15.0 | 15.0 |
| Flaxseed oil | - | - |

### Table S4: Nutritional composition of SFA and PUFA diets

| Ingredient (g/kg) | Saturated fatty acids | Polyunsaturated fatty acids |
|---|---|---|
| Casein | 140.0 | 140.0 |
| L-Cystine | 1.8 | 1.8 |
| Corn starch | 338.552 | 338.552 |
| Maltodextrin | 155.0 | 155.0 |
| Sucrose | 100.0 | 100.0 |
| Olive oil | - | - |
| Soybean oil | - | - |
| Corn oil | - | - |
| Lard | - | - |
| Beef tallow | - | - |
| Anhydrous milkfat | 161.85 | - |
| Cholesterol | - | - |
| Cellulose | 50.0 | 50.0 |
| Mineral mix, nTWD | - | - |
| Sodium chloride | - | - |

| | | |
|---|---|---|
| **Vitamin mix, nTWD** | - | - |
| **Choline bitartate** | 2.75 | 2.75 |
| **Vitamin K1, phylloquinone** | 0.002 | 0.002 |
| **Thiamine (81%)** | 0.013 | 0.013 |
| **TBHQ, antioxidant** | 0.033 | 0.033 |
| **Mineral mix, AIN93-M-MX** | 35.0 | 35.0 |
| **Vitamin mix, AIN93-VX** | 15.0 | 15.0 |
| **Flaxseed oil** | - | 161.85 |
